# A Multiple-trait Bayesian Variable Selection Regression Method for Integrating Phenotypic Causal Networks in Genome-Wide Association Studies

**DOI:** 10.1101/847285

**Authors:** Zigui Wang, Deborah Chapman, Gota Morota, Hao Cheng

**Author notes:** Corresponding author: 2139 Meyer Hall, Department of Animal Science, University of California Davis, 1 Shield Ave., Davis, CA 95616.

## Abstract

Bayesian regression methods that incorporate different mixture priors for marker effects are used in multi-trait genomic prediction. These methods can also be extended to genome-wide association studies (GWAS). In multiple-trait GWAS, incorporating the underlying causal structures among traits is essential for comprehensively understanding the relationship between genotypes and traits of interest. Therefore, we develop a GWAS methodology, SEM-BayesCΠ, which, by applying the structural equation model (SEM), can be used to incorporate causal structures into a multi-trait Bayesian regression method using mixture priors. The performance of SEM-BayesCΠ was demonstrated by comparing its GWAS results with those from multi-trait BayesCΠ. Through the inductive causation (IC) algorithm, three potential causal structures were inferred of 0.9 highest posterior density (HPD) interval. SEM-BayesCΠ provides a more comprehensive understanding of the genotype-phenotype mapping than multi-trait BayesCΠ by performing GWAS based on indirect, direct and overall marker effects. The software tool JWAS offers open-source routines to perform these analyses.

## INTRODUCTION

Genome-wide association studies (GWAS) are widely used to identify associations between single nucleotide polymorphisms (SNPs) and phenotypes (Ozaki *et al.* 2002; Visscher *et al.* 2017; McCarthy *et al.* 2008; Cantor *et al.* 2010). GWAS have successfully mapped quantitative trait loci (QTL) associated with traits of interest, e.g., meat quality and quantity in livestock (Sharma *et al.* 2015), crop yields in plants (Liu and Yan 2018), and diseases in humans (Visscher *et al.* 2017). GWAS are typically based on using linear mixed models to fit one SNP at a time to a single trait (Hackinger and Zeggini 2017). While this allows for a relatively simple statistical model, the interwoven nature of gene expression translates to many traits being correlated with each other (Sodini *et al.* 2018). These correlations can be utilized in multi-trait linear mixed models for GWAS to reduce false positives and increase the statistical power for association mapping (O’Reilly *et al.* 2012; Korte *et al.* 2012).

Conventional multi-trait linear mixed models do not consider the causal relationships between traits. To address this issue, researchers have proposed refining the multi-trait methods with structural equation models (SEM) introduced by Wright (1934) that consider the causal relationship among traits. A model that incorporates causal structures should better reflect underlying genetic mechanisms. Gianola and Sorensen (2004) used SEM to extend conventional multi-trait linear mixed models to accommodate for recursive and simultaneous relationships among traits, which allows traits to be explanatory variables for other traits. Recently, Momen *et al.* (2018, 2019) proposed the SEM-based GWAS (SEM-GWAS) methodology by applying SEM to linear mixed models for GWAS. They showed that while conventional GWAS methodology only provides overall SNP effects, SEM-GWAS can capture the complex causal relationships among traits and further decompose the overall SNP effects into direct and indirect effects.

The SEM-GWAS method proposed by Momen *et al.* (2018, 2019) is based on linear mixed models with a fixed substitution effect for the tested SNP and a random effect with covariances defined by a “genomic relationship matrix” computed from genotypes (VanRaden 2008) to account for genetic relatedness. Most GWAS methods based on the “genomic relationship matrix” implicitly assume all markers affect all traits. However, this assumption is not biologically meaningful, especially in multi-trait analyses involving many traits. Cheng *et al.* (2018b) proposed a general class of multi-trait Bayesian variable selection regression methods that use a broad range of mixture priors, e.g., multi-trait BayesCΠ, where each locus can affect any combination of traits, which allows us to more closely model the true biological mechanisms, e.g, pleiotropy.

The primary goal of this current research is to develop a multi-trait Bayesian regression GWAS method that more closely resembles the underlying biological mechanisms including pleiotropy and causal structure among traits. In this paper, we develop and implement a new GWAS method called SEM-BayesCΠ, which integrates SEM to the multi-trait BayesCΠ, to incorporate the underlying biological mechanism. The performance of SEM-BayesCΠ is studied using real rice data.

## MATERIALS AND METHODS

### Multi-trait Bayesian regression model using mixture priors

Assuming that individuals have all traits measured with a general mean as the only fixed effect, we write the multi-trait model for individual *i* from *n* genotyped individuals as:

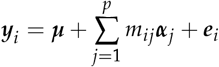

where ***y**_i_* is the vector of phenotypes of *t* traits for individual *i*, ***μ*** is a vector of overall means for *t* traits, *p* is the number of genotyped loci, *m_ij_* is the genotype covariate at locus *j* for individual *i* (coded as 0,1,2), ***α**_j_* is the vector of marker effects of *t* traits for locus *j*, and ***e**_i_* is the vector of residuals of *t* traits for individual *i*. The fixed effects are assigned flat priors. The residuals, ***e**_i_*, are a priori assumed to be independently and identically distributed multivariate normal vectors with null mean and covariance matrix ***R***, which is assumed to have an inverse Wishart prior distribution, *W*^−1^(***S**_e_, ν_e_*).

Allowing each locus to affect any combination of traits, in a multiple-trait Bayesian variable selection method, i.e., multi-trait BayesCΠ (Cheng *et al.* 2018b), the vector of marker effects at locus *j* can be written as ***α**_j_* = ***D**_j_**β**_j_*, where ***D**_j_* is a diagonal matrix whose diagonal element is ***δ**_j_* = (*δ*_*j*1_, *δ*_*j*2_…, *δ_jt_*), where *δ_jk_* is the indicator variable indicating whether the marker effect of locus *j* for trait *k* is zero or not, and ***β**_j_* is a priori assumed to be independently and identically distributed multivariate normal vectors with null mean and covariance matrix ***G***, which is assumed to have an inverse Wishart prior distribution, 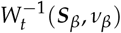. Given that a locus can have an effect on any combination of traits, we use numeric labels “1”, “2”,⋯, “*l*” to represent all 2^*t*^ possible combinations for *δ_j_*, in which case the prior distribution for *δ_j_* is:

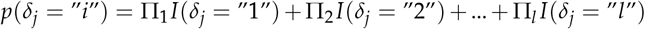

where Π*_i_* is the prior probability that the vector *δ_j_* corresponds to the vector labeled “*i*” and 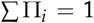. We assume the prior for Π = (Π_1_, Π_2_,…Π_*l*_) is a uniform distribution.

### Structural Equation Model

The linear SEM is composed of two parts: the measurement equation analyzing the relationship between the observable variables and latent variables, and the structural equation capturing the connections among latent variables (Anderson and Gerbing 1988). These two equations can be written as:

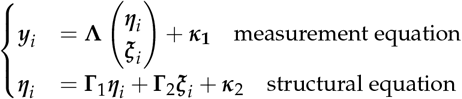

where ***y**_i_* is the vector of observable variables for individual *i*, ***η**_i_* is a *q* × 1 vector of exogenous latent variables, ***ξ**_i_* is a *r* × 1 vector with endogenous latent variables, **Λ** is a *t* × (*q* + *r*) matrix of unknown structural coefficients, ***κ***_1_ and ***κ***_2_ are *t* × 1 vectors of residuals. The details of parameter estimation are discussed in Song and Lee (2012).

In our study, no latent variables are assumed and the sole observable variables are phenotypes. Thus only the causal relationship among observable variables, i.e., phenotypes, are fitted in the SEM model (also known as path analysis (Wright 1921)) as:

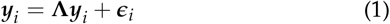

where ***y**_i_* and **Λ** are defined as above, ***ϵ**_i_* represents everything that is not explained by **Λ*y***_*i*_, and **Λ** is an *t* × *t* matrix of structural coefficients representing the causal structure recovered from the Inductive Causation (IC) algorithm as described in the next section.

To illustrate, we assume that the phenotypes of three traits for each individual (i.e., *y*_1_, *y*_2_, and *y*_3_ for traits 1, 2, and 3) have the following causal relationship:

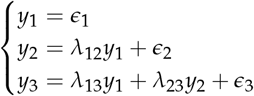

where causal coefficient *λ_ij_* represents that a 1-unit increase in trait *i* results in a *λ_ij_* unit increase in trait *j*. Given the causal structure above, the **Λ** can be written as:

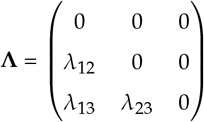

### Searching Causal Structure

As described above, fitting the SEM requires the causal structure among all traits to be known before analysis. To explore the wide-range of possible underlying causal structures, we use the method from Valente *et al.* (2010) to discern the causal structure based on the posterior distribution of residual covariance matrix. The reason we do not directly apply this method to phenotype data is that the covariance among phenotypes is likely confounded by genetic effects. The process of inferring causal structure is composed of three steps:

1. Fit the multi-trait BayesianCΠ model and obtain the posterior distribution of the residual covariance matrix.
2. Follow Valente *et al.* (2010) to derive the conditional independence relationship among traits based on the posterior distribution of the residual covariance matrix. In detail, we derive the residual partial correlation p(*y_i_, y_j_*|h), where h is a set of traits, to test whether trait *y_i_* is conditional independent from *y_j_*. The highest posterior density (HPD) interval of 0.9 was used to make statistical decision. If HPD interval of p(*y_i_, y_j_*|h) contains zero, *y_i_* and *y_j_* are regarded as conditionally independent on h.
3. Apply the IC algorithm (Pearl 2009) to the conditional independence relationship from step 2 to obtain the causal structure as described in the next section.

### Inductive Causation Algorithm

We use the IC algorithm (Pearl 2009) to recover the causal structure among a set of traits, denoted as *U* below, from the conditional independence relationship. The IC algorithm is composed of three steps (Valente *et al.* 2010; Chicharro and Panzeri 2014):

1. For each pair of traits *X* and *Y*, search for a set *S_XY_* ⊆ *U* such that *X*⊥*Y*|*S_XY_* holds. That is, *X* and *Y* are independent, conditional on *S_XY_*. If there is no such *S_XY_*, place an undirected edge between these two traits.
2. If this pair of traits *X* and *Y* are non-adjacent (i.e., no undirected edge between *X* and *Y*) with a common neighbor *Z* (i.e., Z is adjacent to X as well as to Y), and *Z* ∉ *S_XY_*, place arrowheads pointing to Z, i.e., *X* → *Z* ← *Y*.
3. In the partially-oriented graph from step 2, orient as many edges as possible following two requirements:

a. Any alternative orientation will not yield a new *V* structure (i.e., X→Z←Y).
b. Any alternative orientation will not yield a directed cycle.

In summary, we find all the pairs of variables that have a dependent relationship to reconstruct the basic structure of the underlying causal network in step 1. Then we find all the *V* structures in the network in step 2 and prevent the creation of new V structures or directed cycles in step 3.

### SEM-BayesCΠ

Assume 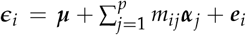 in equation (1) and follow assumptions in multi-trait BayesCΠ, the SEM-Bayes*C*Π model can be written as:

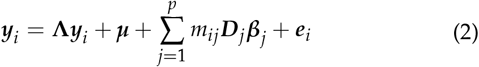

Move **Λ*y***_*i*_ from the right side to the left side of equation (2), and define **Λ**^*^ = ***I*** − **Λ**, where ***I*** is a *t* × *t* identity matrix and **Λ** is a *t* × *t* matrix of structural coefficients based on the discerned causal structure, the model becomes:

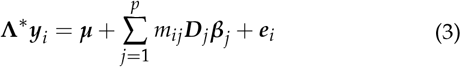

To guarantee that the structural coefficient is identifiable, we assume that the residuals for each trait of individual *i* are independent with each other, which means the residual covariance matrix is diagonal (Wu *et al.* 2010; Momen *et al.* 2018). The vector of all non-zero elements in **Λ**, e.g., ***λ*** = [*λ*_12_, *λ*_13_, *λ*_23_], is assumed to have a prior distribution:

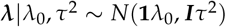

where **1** is a vector of ones, ***I*** is the identity matrix, and *λ*_0_ is a known mean for all elements in ***λ***. *τ*^2^ is a tuning parameter to adjust the sharpness degree of the prior (Gianola and Sorensen 2004). In this paper, we set *λ*_0_ = 0 and *τ*^2^ = 1. The priors for remaining parameters are the same as in the section Multi-trait Bayesian regression model using mixture priors.

Gibbs samplers are used to draw samples for all parameters. The full conditional distribution to draw samples for ***λ*** is shown below. The derivations of the full conditional distributions of the remaining parameters of interest for Gibbs samplers are in Cheng *et al.* (2018b).

#### Full conditional distribution of Λ

We follow Gianola and Sorensen (2004) to obtain the full conditional distribution of **Λ**, with the difference between our derivation and Gianola and Sorensen (2004) being that we specify the causal structure with positions of parameters in the **Λ**. Let Ω denote all parameters except ***λ*** in the SEM-BayesCΠ and use the causal structure 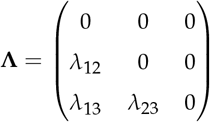 as an example, the left hand side of equation (3), **Λ**^*^***y**_i_*, can be written as:

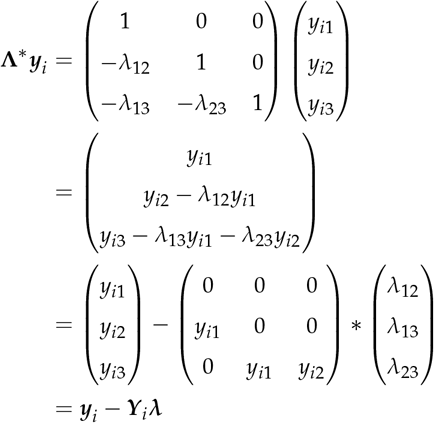

The conditional posterior distribution of ***λ*** can be written as:

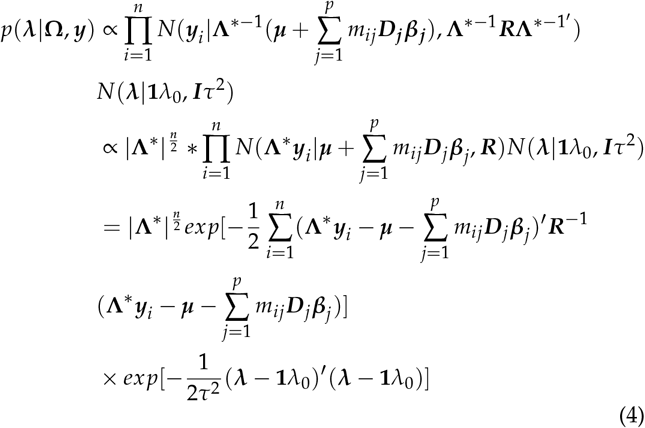

Setting 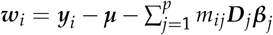, equation (4) can be written as:

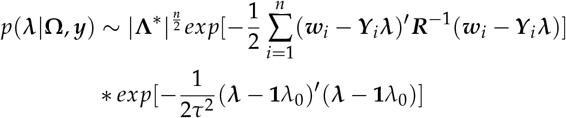

Following the derivation in Gianola and Sorensen (2004) and the fact that |**Λ**^*^| = 1 in a recursive system, the full conditional distribution of ***λ*** is

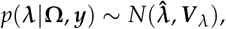

where

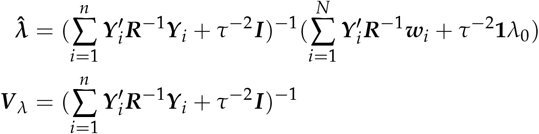

### Decomposition of SNP Effects

In SEM-BayesCΠ, the marker effect for locus *j*, ***α**_j_*, is considered as the vector of direct marker effects of *t* traits. The indirect effect of locus *j* of *t* traits can be calculated as 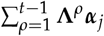. The overall effect of locus *j* on *t* traits is computed as 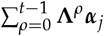 or (***I*** − **Λ**)^−1^ ***α**_j_*, which is the summation of both direct and indirect effects of locus *j*. For example, given a causal structure 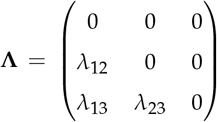, the direct effect for locus *j* on three traits is 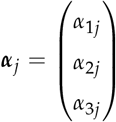, and the indirect effect for locus *j* on three traits is calculated as 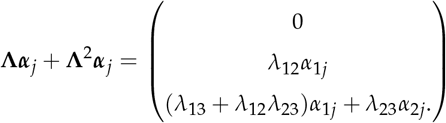, and the overall effect of locus *j* on trait *k* is 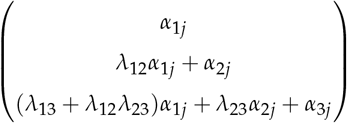.

### Inference of Association based on Genomic Windows

Markers in a genomic window are usually highly correlated, indicating that any single marker may not show a strong association with the trait even though a causal variant exists in the window. In this paper, we make an inference of association based on genomic windows, because multiple markers inside a genomic window may jointly capture the signal from the causal variant (Fernando and Garrick 2013; Fernando *et al.* 2017).

To make an inference of association based on genomic windows, posterior distribution for the proportion of the genetic variance explained by markers in genomic window *w*, *q_w_*, is estimated from MCMC samples of overall, direct, and indirect marker effects as follows. For one MCMC sample of all marker effects on one trait, let ***α**_direct_*, ***α**_indirect_*, and ***α**_overall_* denote direct, indirect, and overall effects of all markers respectively.

The genetic value that is attributed to genomic window *w* is calculated as:

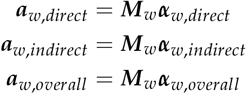

where ***M**_w_* is a matrix of marker covariates in window *w* and ***α**_w,direct_*, ***α**_w,indirect_*, and ***α**_w,overall_* are the MCMC samples of direct, indirect, and overall marker effects for SNPs in window *w*. Then the variance explained by the genomic window *w* is defined as:

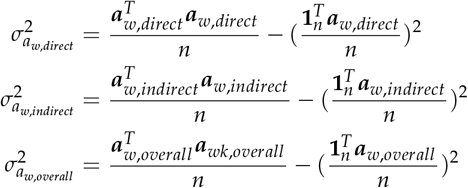

Similarly, the total genetic variance is calculated as:

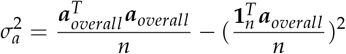

The proportion of the genetic variance explained by direct, indirect, and overall marker effects in the genomic window *w* is calculated as:

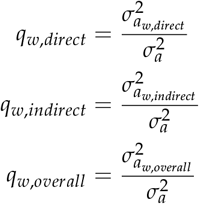

Given the MCMC samples of *q_w_*, the window posterior probability of association (WPPA) is calculated as the proportion of MCMC samples of *q_w_* that exceed a specific value *T* (Fernando and Garrick 2013; Chen *et al.* 2017; Lloyd-Jones *et al.* 2017). In this paper, associations are tested for non-overlapping windows of 100 SNPs, and genomic windows that explain over 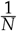 of the total genetic variance were deemed to be of potential interest (i.e., 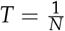, where *N* is the total number of windows).

### Threshold of WPPA

It has been shown that using a threshold of WPPA = *α* to declare a significant genomic window restricts the proportion of false positives (PFP) to < 1 − *α* (Fernando *et al.* 2017). A previous GWAS (Zhao *et al.* 2011) identified significantly associated SNPs in chromosome 6 for flowering time in Arkansas (FTA) using the same dataset. A threshold of WPPA = 0.8 in our GWAS analysis resulted in similarly significant signals. This result suggests that a WPPA of 0.8 is reasonable for declaring a significant genomic window.

### Data Analysis

The Rice Diversity Panel with 413 Oryza sativa individual accessions was used in the analysis. Three traits were considered, including plant height (PH), flowering time in Arkansas (FTA), and panicle number per plant (PN) in our GWAS. After removing the records with missing data for these three traits, 370 individuals with 36,901 SNPs genotyped were included in our analysis. The phenotypic and genotypic data were publicly available for download from http://www.ricediversity.org/.

We have implemented SEM-BayesCΠ in JWAS (Cheng *et al.* 2018a), an open-source, publicly available package for single-trait and multi-trait whole-genome analyses written in the freely available Julia language.

## RESULTS

### Causal Structure and Structural Coefficients

The causal structure among three traits is inferred by the IC algorithm from the estimated posterior distribution of the residual covariance matrix in the multi-trait BayesCΠ model. Figure 1 shows three potential phenotypic causal structures among traits PH (*y*_1_), FTA (*y*_2_), and PN (*y*_3_) recovered for the 0.9 HPD interval. The causal structure matrices for IC1 (**Λ**_1_), IC2 (**Λ**_2_), and IC3 (**Λ**_3_) are:

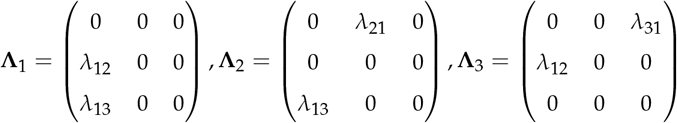

**Figure 1.**
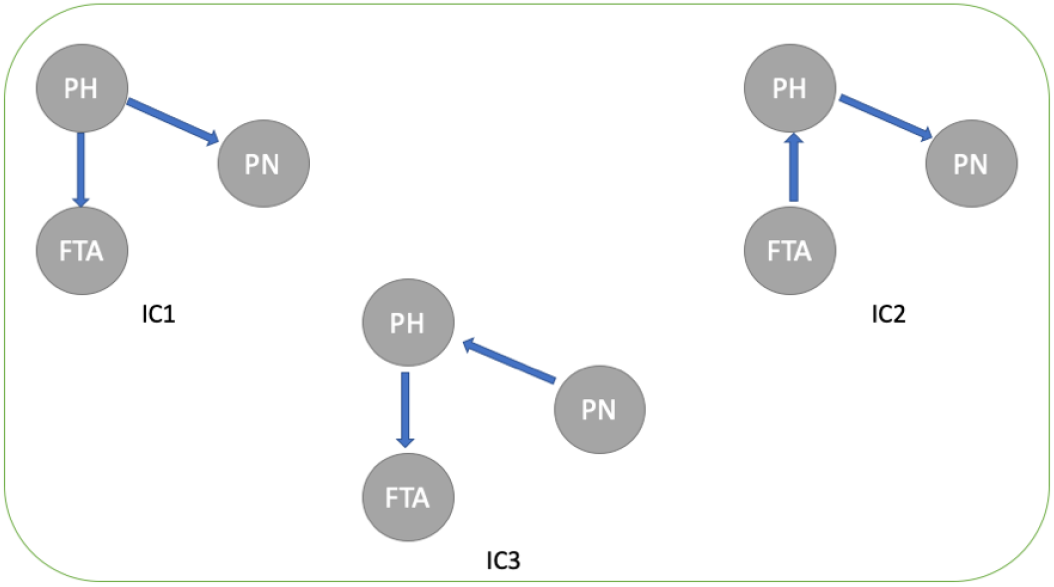
Causal structures among plant height (PH), flowering time in Arkansas (FTA), and panicle number per plant (PN) inferred from the IC algorithms. The edges connecting two traits represent non-null partial correlations as indicated by 0.9 HPD interval. The arrows represent the direction of causal effects.

These three causal structures are fitted in the SEM-BayesCΠ model, and samples from posterior distributions for coefficients in these causal structures are obtained. The 90% credible intervals for structural coefficients in IC1, IC2, and IC3 are shown in Table 1. Structure coefficients indicate the strength of causal relationship between traits. Though all credible intervals shown in Table 1 do not include zero, those for *λ_PH→PN_* and *λ_PN→PH_* in all three causal structures are close to zero, indicating that these two causal relationships might be weak.

**Table 1.** The 90% credible interval for causal structural coefficients in the three causal structures

| $\lambda$ | IC1 | IC2 | IC3 |
| --- | --- | --- | --- |
| $\lambda_{PH \rightarrow FTA}$ | (0.29, 0.56) | NA | (0.31, 0.53) |
| $\lambda_{PH \rightarrow PN}$ | (-0.22, -0.05) | (-0.20, -0.04) | NA |
| $\lambda_{FTA \rightarrow PH}$ | NA | (0.25, 0.42) | NA |
| $\lambda_{PN \rightarrow PH}$ | NA | NA | (-0.26, -0.08) |

PH, FTA, and PN represent traits plant height, flowering time in Arkansas, panicle number per plant, respectively. $\lambda_{a \rightarrow b}$ represents the causal effects of trait *a* on trait *b*. NA denotes structural coefficients those do not exist in the causal structure.

### Decomposition of SNP Effects in SEM-BayesCΠ

The causal structure IC1 was also inferred by Tabu, Hill Climbing, General 2-Phase Restricted Maximization and Max-Min Hill Climbing algorithm in Yu *et al.* (2019) using a similar dataset. Thus, for the simplicity of our presentation, but without loss of generality, IC1 is used in this section to demonstrate the performance of SEM-BayesCΠ in capturing the causal relationship among traits and decomposing the total SNP effects. Direct, indirect, and overall SNP effects for all markers are estimated from SEM-BayesCΠ.

#### Direct SNP Effects

In SEM-BayesCΠ, direct SNP effects are assigned mixture priors, where each locus can affect any combinations of traits directly. As shown in Figure 2, the posterior distribution of the parameter Π is obtained, and markers show different levels of pleiotropy for direct SNP effects.

**Figure 2.**
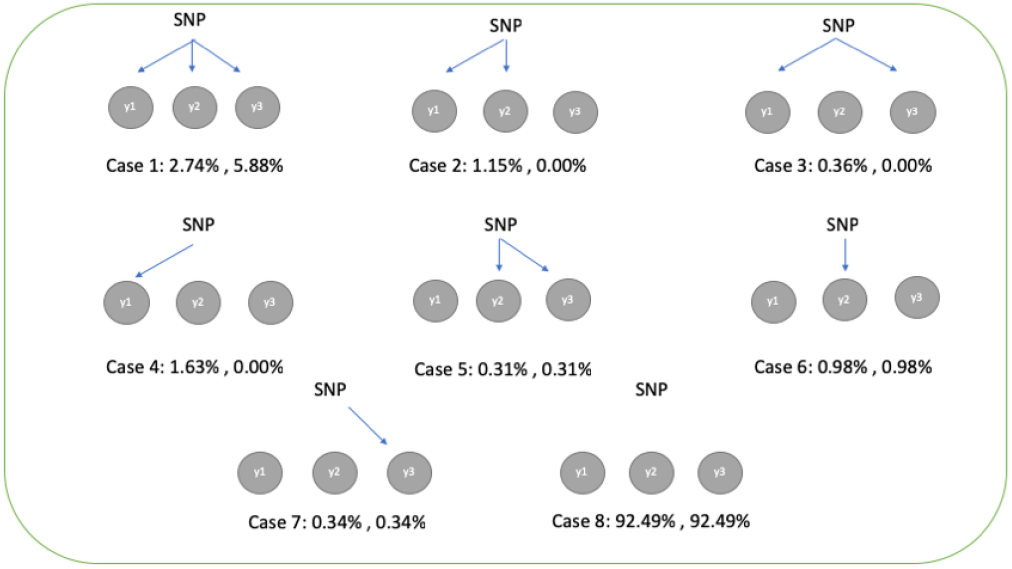
Estimated proportion of markers affecting combinations of traits, Π, from SEM-BayesCΠ incorporating IC1 causal structure. For each scenario, it is estimated for both direct SNP effects (left) and overall SNP effects (right)

#### Indirect SNP Effects

In SEM-BayesCΠ, samples from posterior distributions of indirect effects are obtained using joint samples from posterior distributions of **Λ** and direct SNP effects ***α**_j_*.

#### Overall SNP Effects

In IC1, for trait PH, the overall SNP effect is equal to the direct SNP effect, because there is no intermediate trait. For trait FTA, the overall SNP effect is composed of direct SNP effect and indirect SNP effect transmitted from PH. So the overall SNP effect for FTA is given by summing the direct SNP effect and indirect SNP effect. Similarly, for trait PN, the overall SNP effect is obtained by summing the direct SNP effect and indirect SNP effect transmitted from PH.

In SEM-BayesCΠ, each SNP can have direct effects on any combination of traits, and the parameter Π is used to estimate the proportion of SNPs having different levels of pleiotropy. Indirect SNP effects on one trait are transmitted from direct SNP effects on other intermediate traits. For example, in causal structure IC1, the indirect SNP effects on FTA is transmitted from direct SNP effects on the trait PH. A SNP having no direct effect on FTA may have overall effects on FTA if its direct effect on trait PH is non-zero. The proportions of markers affecting different combinations of traits through overall SNP effects can also be estimated. This result is shown in Figure 2, and different probabilities are observed for some cases between overall and direct SNP effects. More SNPs have effects on all traits simultaneously when overall SNP effects are considered compared to direct SNP effects (case 1). If only overall SNP effects are considered, some cases having non-zero probabilities for indirect SNP effects are hidden by the causal relationships among traits (cases 2-4). The same patterns for direct and overall SNP effects are observed in cases 5-8 because there is no causal relationship between trait FTA and PN.

### GWAS Result

The results of GWAS from multi-trait BayesCΠ and SEM-BayesCΠ incorporating three causal structures are shown in Figure 3. Significant signals are found only for trait FTA, and results of GWAS for this trait from multi-trait BayesCΠ and SEM-BayesCΠ incorporating three causal structures are shown in Figure 3. A threshold of WPPA = 0.8 is adopted to declare a significant genomic window. The blue point represents genomic window A containing SNPs from “id5012741” to “id5013321” on chromosome 5. The red point represents window B containing SNPs from “id6005556” to “id6006216” on chromosome 6. In multi-trait BayesCΠ, window B achieved WPPA 0.90. In multi-trait SEM-BayesCΠ incorporating IC1 causal structure, window A achieved WPPA 0.94, and genomic window B achieved WPPA 0.84. In multi-trait SEM-BayesCΠ incorporating IC2 causal structure, only window B has significant signals and achieved WPPA 0.92. In SEM-BayesCΠ incorporating IC3 causal structure, window B achieves WPPA 0.94, and window A achieves WPPA 0.84. Though window A and B are not found as significant in all methods using a threshold WPPA = 0.8, peaks are always observed for window A and B in all methods with a WPPA at least around 0.8. The same peak on chromosome 6 was also identified in (Zhao *et al.* 2011) using a similar dataset.

**Figure 3.**
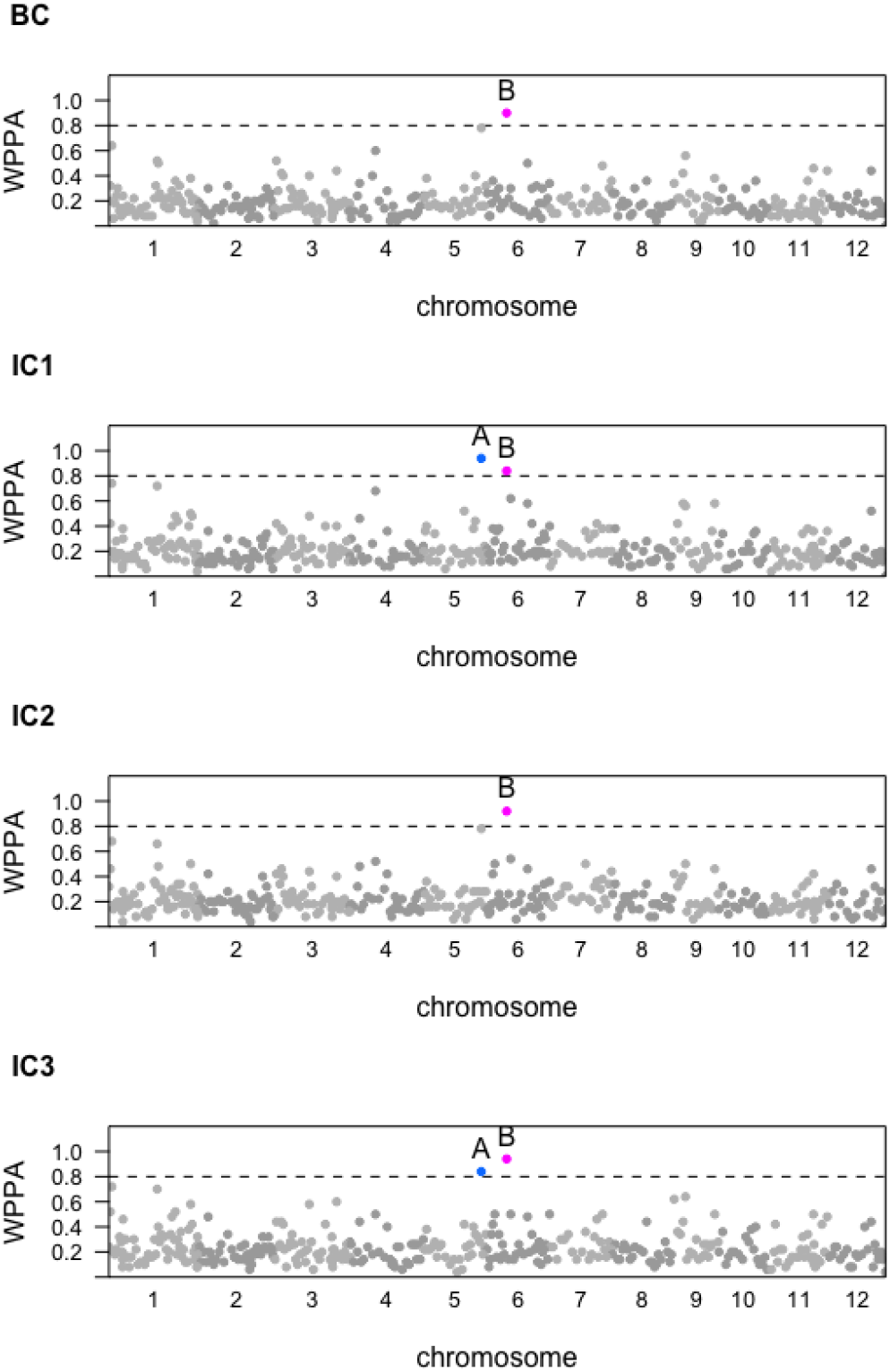
GWAS results from multi-trait BayesCΠ (BC) and SEM-BayesCΠ (IC1, IC2, and IC3) incorporating three causal structures based on the overall SNP effects for the trait flowering time at Arkansas(FTA). X-axis represents the locations for genomic windows along the 12 chromosomes; Y-axis represents window posterior probability of association (WPPA). Colored points represent genomic windows with WPPA >0.8.

The overall SNP effects are partitioned into direct and indirect effects, and GWAS are performed for the direct, indirect, and overall SNP effects separately for trait FTA. The results of GWAS for trait FTA are shown in Figure 4. A threshold of WPPA = 0.8 is adopted to declare a significant genomic window. In Figure 4, the blue point represents a window containing SNPs from “id5012741” to “id5013321” on chromosome 5, and the red point represents a window containing SNPs from “id6005556” to “id6006216” on chromosome 6. In the overall effects from SEM-BayesCΠ, window A achieved WPPA 0.94, and window B achieved WPPA 0.84. In the direct effects from SEM-BayesCΠ, window A achieves WPPA 0.90, and window B achieves WPPA 0.84. In indirect effects, no window is identified as significant, though peaks on chromosome 5 and 6 are also observed. WPPA for direct SNP effects are more correlated with those for overall SNP effects. Further, magnitudes for overall, direct, and indirect SNP effects are also shown in Figure 5 (Appendix). Though most large overall SNP effects consist of a large direct SNP effect and a relatively small indirect SNP effect, the indirect effects of some SNPs play an important role, e.g., SNP “id1020584”, as shown in Figure 5, showed direct and indirect SNP effects of opposite magnitudes, resulting in overall SNP effects of about zero.

**Figure 4.**
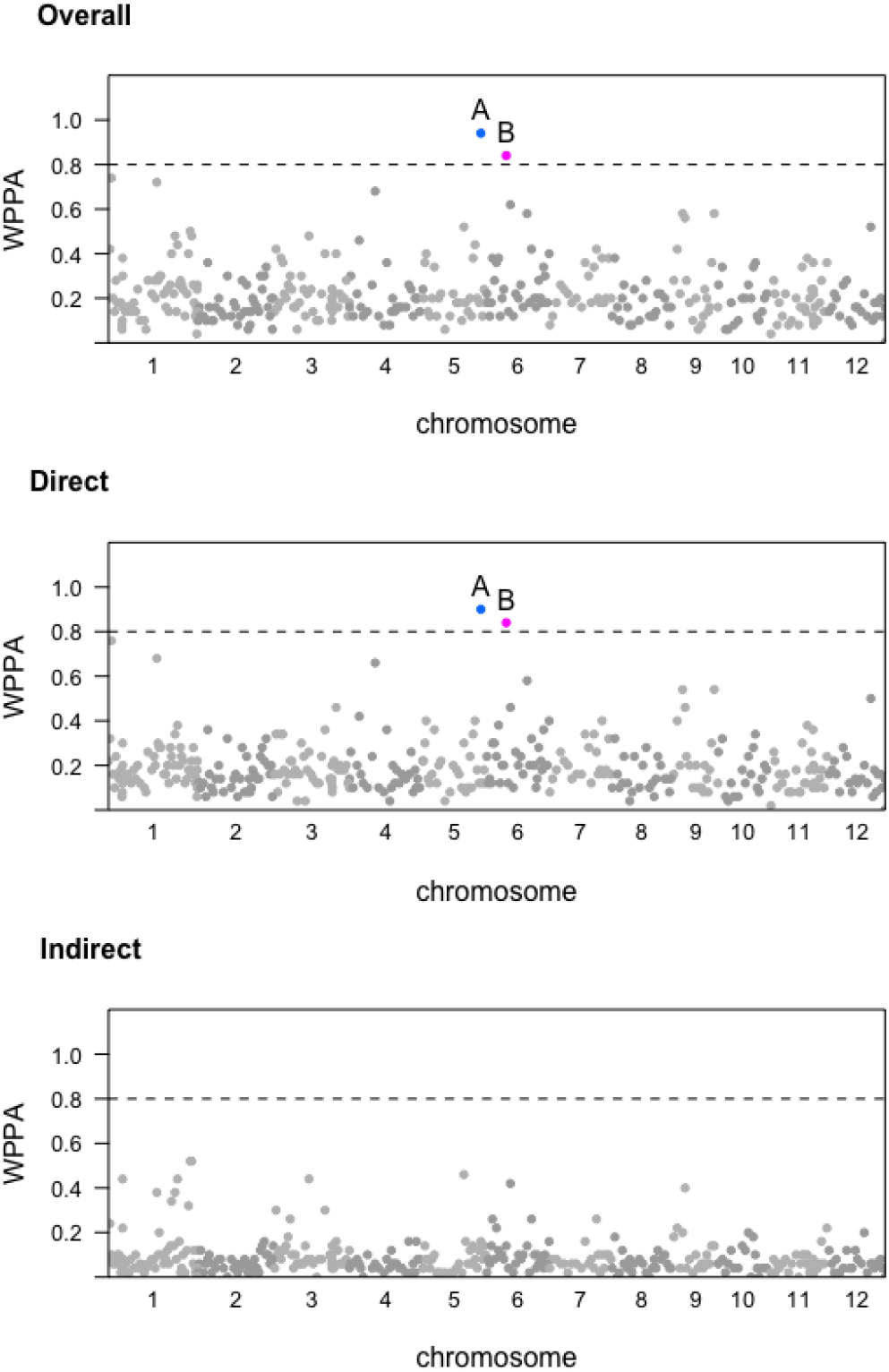
GWAS results based on direct, indirect and overall SNP effects from SEM-BayesCΠ incorporating IC1 causal structure for the trait flowering time at Arkansas (FTA). The horizontal dash line represents the threshold 0.8. X-axis represents the location of genomic windows along the 12 chromosomes; Y-axis represents window posterior probability of association (WPPA). Colored points represent genomic windows with WPPA > 0.8.

## DISCUSSION

The complex causal relationships among multiple traits are usually not considered in conventional multi-trait GWAS. Here we propose the SEM-BayesCΠ method to incorporate pre-inferred causal structures among multiple traits into a multi-trait Bayesian regression method using mixture priors. SEM-BayesCΠ accounts for causal structures among traits, and has the potential advantage of estimating causal effects, as well as providing a comprehensive understanding of the underlying biological mechanism.

### Causal Structure

The causal structure is assumed to be known in SEM-BayesCΠ, and it is usually discerned by three types of algorithms: the constraint-based algorithm, the score-based algorithms, and the hybrid algorithms. The IC algorithm (Pearl 2009; Valente *et al.* 2010) discussed in this paper is a typical constraint-based algorithm, which is based on conditional independence tests. The score-based algorithms apply the heuristic optimization techniques, which set an initial graph structure and assign an initial goodness-of-fit score to it, and then maximize the goodness-of-fit score to obtain the most possible causal structure. The hybrid algorithm is a hybrid of both the constraint-based and the score-based algorithms. It utilizes conditional independence tests to reduce the space of candidate causal structures, and uses network scores to identify the optimal structure among them (Scutari 2014). The causal structures inferred from these algorithms may be different. Note that different evaluation criteria may also result in different outcome causal structures. For example, in this paper, if we choose 0.99 instead of 0.9 HPD interval to search for causal structures, there will be no edge between the traits PH and PN.

### Decomposition of SNP Effects

In some previous analysis (Mi *et al.* 2010; Momen *et al.* 2018, 2019), the indirect SNP effect of locus *j* of *t* traits is obtained by multiplying the estimated **Λ**, 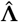, and estimated direct SNP effects, 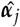, as 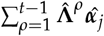. This is similar to using posterior means of causal structural coefficients and direct SNP effects for calculation of the indirect SNP effects. In our method, indirect SNP effects are estimated using joint samples from posterior distributions of **Λ** and ***í**_j_*. We compared these two approaches for indirect SNP effect estimation, and found that the indirect effects estimated from these two approaches are slightly different. The SEM-BayesCΠ approach should be used in indirect SNP effect estimation due to the fact that **Λ** and ***α**_j_* may be highly dependent.

### GWAS

Figure 3 shows similar GWAS results from the SEM-BayesCΠ and BayesCΠ. This is reasonable since the SEM-BayesCΠ model can be reduced to a model similar to BayesCΠ by reparameterization, indicating that the joint likelihood functions of SEM-BayesCΠ and BayesCΠ are similar. The agreement between SEM-BayesCΠ and BayesCΠ results can be considered as verification that SEM-BayesCΠ gives the correct result in GWAS. As shown in Figures 2, 4 and 5, SEM-BayesCΠ provides a more comprehensive understanding of the underlying biological mechanism.

**Figure 5.**
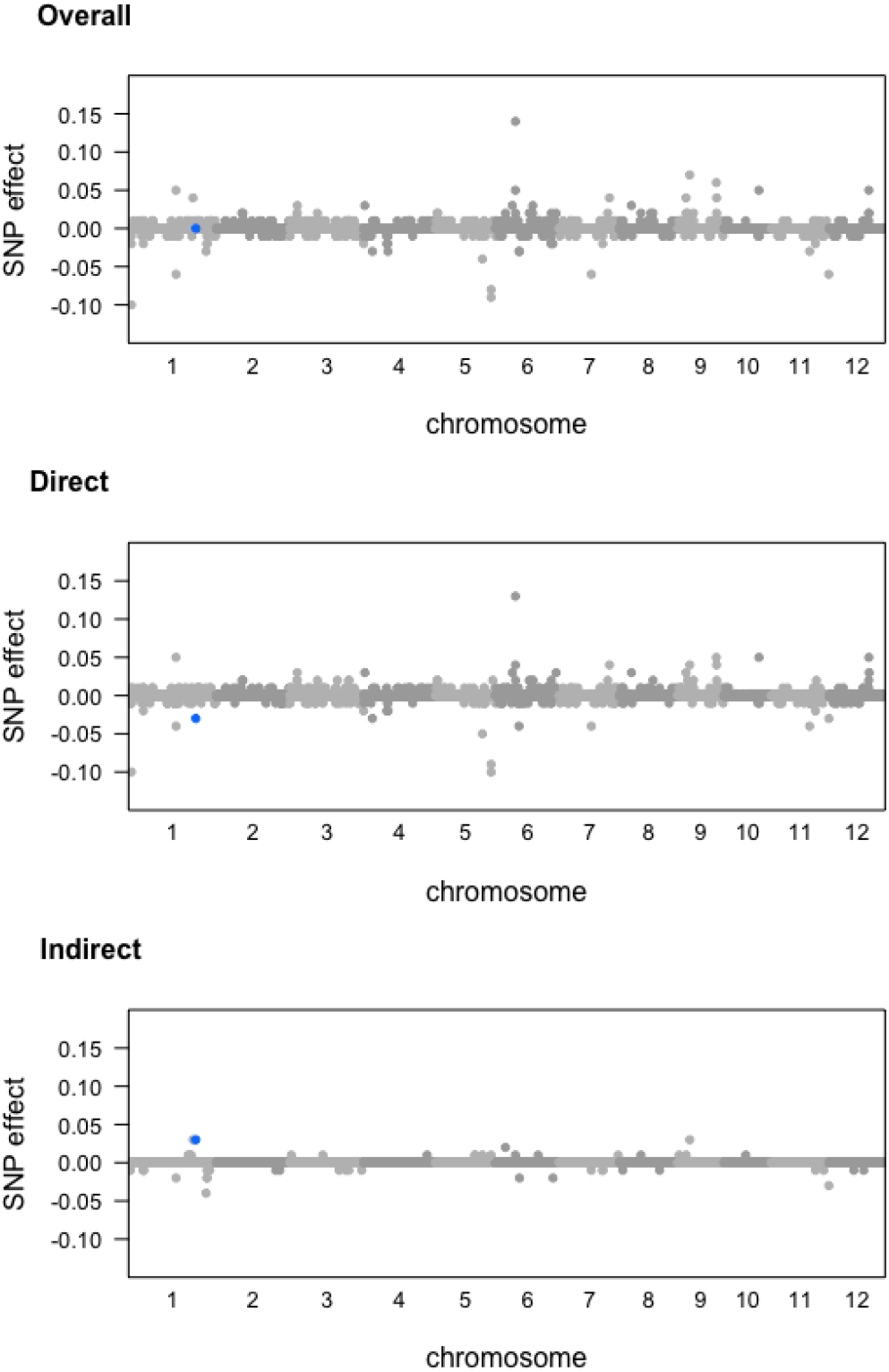
Magnitude of direct, indirect and overall SNP effects from SEM-BayesCΠ incorporating IC1 causal structure for the trait flowering time at Arkansas (FTA). X-axis represents the location of SNPs along the 12 chromosomes. Y-axis represents the magnitude of the marker effects. The blue points represents the SNP “id1020584”.

### Conclusion

SEM-BayesCΠ provides more interpretation into biological mechanisms by decomposing the overall SNP effects into direct and indirect SNP effects. In SEM-BayesCΠ, posterior distributions of the overall, direct, and indirect SNP effects, as well as causal structure coefficients, are obtained, which are used to make inferences about these parameters. In our genomic window-based GWAS, the posterior distributions are obtained for the proportion of variance attributed to a genomic region to detect causal loci (i.e., the use of WPPA). The level of gene pleiotropy, e.g., proportion of markers affecting different combinations of traits as shown in Figure 2, can also be further dissected into direct and indirect SNP effects. In summary, compared to conventional GWAS, SEM-BayesCΠ offers a more comprehensive understanding of the underlying biological mechanisms including pleiotropy and causal relationships among traits.

## ACKNOWLEDGEMENTS

We want to thank Bruno D. Valente for his explanation of the causal structure searching method and Mehdi Momen for his clarificationon the IC algorithm. This work was supported by the United States Department of Agriculture, Agriculture and Food Research Initiative National Institute of Food and Agriculture Competitive grant no. 2018-67015-27957.

## APPENDIX

Estimated direct, indirect, and overall SNP effects from SEM-BayesCΠ incorporating the IC1 causal structure for the trait flowering time at Arkansas (FTA) are shown in Figure 5. The SNP “id1020584” has direct effect −0.031 on trait FTA, while its indirect effect transmitted through PH is 0.034. Thus, the overall effects of SNP “id1020584” is about zero.

